# Polyelectrolyte chain at thermal equilibrium: A comparison between the density functional theory framework (DFT) and the molecular dynamic simulations (MD)

**DOI:** 10.1101/2021.02.17.431088

**Authors:** Mike J. Edwards

**Affiliations:** Leibniz-Institut für Polymerforchung, Hohe Stra*β*e 6, 01069 Dresden, Germany

## Abstract

By means of the density functional theory framework (DFT) as well as the molecular dynamic simulations (MD), a polyelectrolyte chain (PE) in the good solvent conditions at thermal equilibrium is studied. The strength of the electrostatic interactions is varied by the Bjerrum length of the solvent. It turns out that average extension of a PE scales with the degree of polymerization, very much similar to a neutral polymer chain in good solvent. Remarkably, the difference between a PE and a neutral chain appears to be solely in the correlations among monomers which are stored in the Virial coefficients. Interestingly, upon increasing the Bjerrum length of solvent, the chain shrinks. This outcome is confirmed by the DFT framework as well as the MD simulations.

**SIGNIFICANCE:** The significance of this study is that it strongly criticizes the idea (already mentioned in T. Kreer, *Soft Matter*, **12**, 3479 (2016)) that the PEs behave similar to a neutral ideal chain. This study could be useful in our understanding of biopolymers.

## INTRODUCTION

Polymers are formed when a large number of repeating units (monomers) are connected through covalent bonds in which the monomers share their valence electrons (1). Polymers normally consist of 10^4^ to 10^5^ monomers. The Polyelectrolytes (PE) are polymers with charged monomers. The PEs are present in almost all biological systems such as in cells as Glycol, in the synovial joints of mammals as aggregan etc (1). When a PE is placed in a solvent, the oppositely and negatively charged particles called counterions (CI) are dissociated from the PE into solvent to fulfill the electroneutrality principle. The size of the CIs are much smaller than monomers so their excluded volume interactions with monomers could be ignored. This makes methods that consider the CIs implicitly more efficient than the explicit methods. In this study, the solvent and CI molecules are implicitly taken into account. The CIs screen the electrostatic interactions between monomers. So the presence of CIs with moderate concentration turns the long-range electrostatic interactions into short-range interactions. This property makes handling of the electrostatic interactions in the biological systems straightforward. In Fig. (1), a 2D schematic view of a PE and the CIs is indicated.

**Figure 1:**
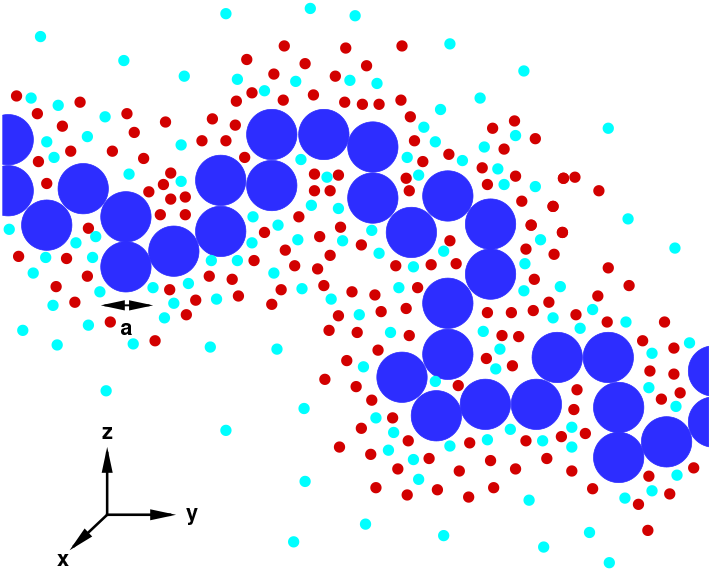
A 2D schematic view of the PE and the CI molecules. The blue shows the PE monomers, the red shows the oppositely charged particles and light blue shows the equally charged particles.

## METHODS

In this study, I employ two absolutely separate methods, the DFT framework and the MD simulations. In the following, I provide short descriptions of these methods.

### Density functional theory framework (DFT)

In many body systems, the golden key is the density distribution function which is a function of the coordinates. Once the density function of a system is known, all physical quantities are specified. To this end, the grand potential functional is written in terms of different Helmholtz free energies in the system plus a constraint to fix the number of particles. Once, the grand potential functional is known, the equilibrium quantities could be calculated through energy minimization principle. This is done by taking variational derivative of the grand potential with respect to density as well as its normal derivative with respect to other quantities. At equilibrium conditions, all these derivatives must vanish. This gives us a set of coupled equations to be solved simultaneously.

In the case of a polymer chain, there are two major effects. Firstly, the entropic elasticity which tries to contract the chain. Secondly, the excluded volume interactions between monomers which tends to stretch the chain. The balance between these forces, holds the chain in a certain average extension. The concept of ideal chain refers to the a chain without the excluded volume interactions. In this case, the density distribution function is Gaussian and the average chain extension is *l* = *aN*^1/2^. The Gaussian properties results from the N-vector model in which a large number of vectors rotate freely in space (2–4). Once, the excluded volume interactions is turned on, the density distribution function becomes Parabolic and the chain extension becomes a function of correlations. To see this, I proceed with the DFT framework (5, 6). The grand potential functional is written as follows,

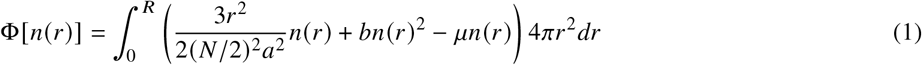

Here, the origin of coordinate is places in the center of the chain and due to the spherical symmetry of the problem, the spherical polar coordinates is chosen. The first term in Eq. (1), refers to the entropic elasticity of *N*/2 monomers. The second term represents the excluded volume interactions among monomers and *b* is the second *Virial* coefficient (2–6). The second Virial coefficient is defined as follows,

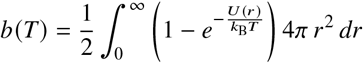

where *U(r)* is the binary potential energy between between monomers. The energy minimization as well as the fixed number of monomers imply the following set of coupled equations to be solved simultaneously,

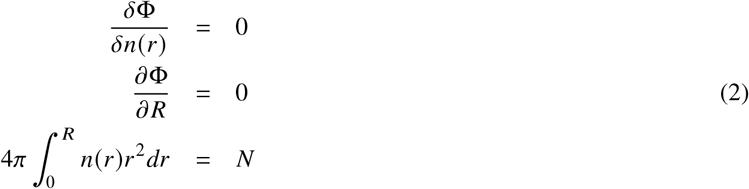

Eq. (2) could be solved analytically and it gives the following solutions,

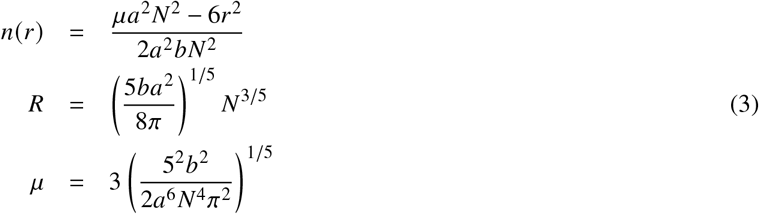

Eq. 3 reveals that the density distribution function is Parabolic. Additionally, it turns out that the chain extension scales as ~ *N*^3/5^ which is slightly larger than the ideal chain model. Remarkably, the chain extension scales as ~ *a*^2/5^. This power law is much lower than ideal chain model which scales linearly with the Kuhn length. Finally, the chain length scales as ~ *b*^1/5^. This power law indicates a weak dependence of the chain length on the correlations between monomers.

A PE consists of charged monomers which interact via electrostatic interactions. Once, a PE is placed into a solvent, the positive and negative counterions dissociate into the solvent as well. The existence of these counterions gives rise to electrostatic screening effect. Hence, in biological systems, the electrostatic interactions are often effective withing the screening length. This means that, despite the fact that the electrostatic interactions are intrinsically long-range, however, in biological medium the they are short-range. The primary competitor of the electrostatic interactions is the thermal energy. The length scale in which the electrostatic interactions balance the thermal energy is called the *Bjerrum* length that is defined as,

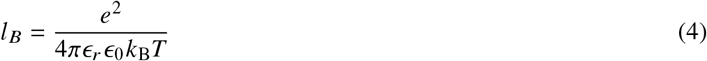

where *e* is the elementary charge, *ϵ*_0_ the vacuum permittivity, *ϵ_r_* the dielectric constant of the solvent, *k*_B_ the Boltzmann constant and *T* the absolute temperature in Kelvins. The water is dominant in biological media. At room temperature *T* = 300*K*, the dielectric constant of water estimated as 80, so the Bjerrum length of water gets *l_B_* = 0.7*a*. The temperature in reduced units is taken as *T* = 1.68 *ϵ/k*_B_.

The second competitor of the electrostatic interactions is the screening. To estimate the length scale in which screening takes place, the Poisson-Boltzmann differential equation needs to be solved subject to appropriate boundary conditions. To do that the Poisson-Boltzmann differential equation is written as follows,

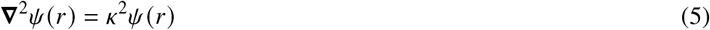

where *ψ*(*r*) is the electrostatic potential scaled by *k*_B_*T/e* and the screening length of the electrostatic interactions is defined as follows,

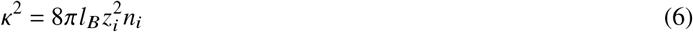

with *z_i_* the valency of counterions and *n_i_* the density of counterions. The solution of Eq. 6 in spherical polar coordinates subject to the boundary conditions that at (*r* → ∞) the potential vanishes and at (*r* ≪ *K*^−1^ the potential behaves like a point charge, is given as follows,

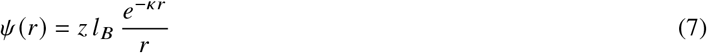

where *z* is the valency of monomers. Eq. (7) is the *Debye-Hückel* electrostatic potential which is widely used in biological systems (7).

### Molecular dynamic simulation (MD)

The most efficient computer simulation method for many body systems is the molecular dynamic simulation (MD). In the MD, the Newton’s equation of motion is solved for each particle. The most efficient discretization scheme for MD is the Leap-Frog method in which the half-step velocities are used to calculate the new positions and velocities (8). The accuracy of Leap-Frog is up to Δ*t*^3^. The coarse-grain model of polymer is constructed by Kremer-Grest model (KG) (9). In the KG model the excluded volume interactions is defined by the Lennard-Jones potential shifted and cut-off as follows,

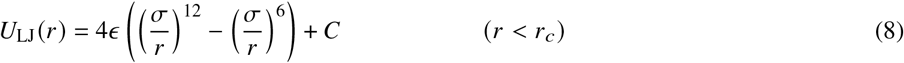

For good solvent conditions (*r*_*c*_ = 2^1/6^ and *C* = *ϵ* and for poor solvent conditions (*r*_*c*_ = 2× 2^1/6^ and *C* = 0.031*ϵ*. The connectivity throughout the chain is constructed via finitely extensible nonlinear elastic potential (FENE) which is given as follows,

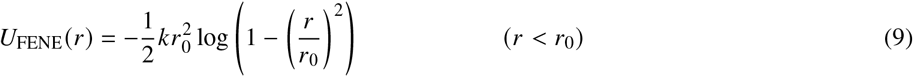

where *k* = 30 and *r*_0_ = 1.5σ. The hydrodynamic drag and the thermal fluctuations are introduced through the *Langevin* thermostat. The Langevin thermostat conssists of two forces which are given as follows,

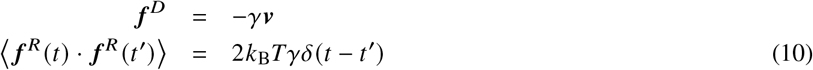

where *γ* is the solvent friction coefficient and *v* is monomer velocity. The electrostatic interactions is defined by the Debye-Hückel potential in Eq. (7). The numerical values of simulation parameters are listed in Table (1).

**Table 1:** The numerical values of the system parameters

|  |  |  |
| --- | --- | --- |
| Temperature | $T$ | $1.68 \epsilon/k_B$ |
| MD simulations time step | $\Delta t$ | $2 \times 10^{-3} (m\sigma^2/\epsilon)^{1/2}$ |
| MD simulations run time | $t$ | $10^7 \Delta t$ |
| Lennard-Jones cutoff for good solvent | $r_c$ | $2^{1/6} \sigma$ |
| Lennard-Jones cutoff for poor solvent | $r_c$ | $2 \times 2^{1/6} \sigma$ |
| FENE maximum extension | $r_0$ | $1.5 \sigma$ |
| FENE elastic constant | $k$ | $30 \epsilon/\sigma^2$ |
| Hydrodynamic friction coefficient | $\gamma$ | $5 (m\epsilon/\sigma^2)^{1/2}$ |
| Bjerrum length | $l_B$ | $0.7 \sigma$ |
| CI concentration | $n_i$ | $10^{-1} \sigma^{-3}$ |
| CI valency | $z_i$ | 1 |
| Monomer valency | $z$ | 1 |
| Debye screening length for $l_B = 0.7 \sigma$ | $\kappa^{-1}$ | $0.75 \sigma$ |
| Debye screening length for $l_B = 2.0 \sigma$ | $\kappa^{-1}$ | $0.45 \sigma$ |
| Kuhn length | $a$ | $0.96 \sigma$ |
| Second Virial coefficient for polymer in good solvent | $b$ | $2.08 \sigma^3$ |
| Second Virial coefficient for PE in good solvent $l_B = 0.7$ | $b$ | $2.99 \sigma^3$ |
| Second Virial coefficient for PE in good solvent $l_B = 2.0$ | $b$ | $2.5 \sigma^3$ |

## RESULTS

Fig. (2) shows the mean squared end-to-end distance 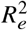 of a variety of chains in terms of degree of polymerization *N*. The plots show a very good agreement between the MD simulations and the DFT framework 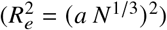 for a neutral chain in poor solvent. This reveals that the chain in poor solvent is completely collapsed. The neutral chain in good solvent stretches to a certain extension and it is fairly scaled as ~ *N*^3/5^. However, the magnitude of the extension is below the prediction of the DFT framework. This could be somewhat natural as the DFT framework assumes that all monomers correlate to each other. While, in the realistic situations, a fraction of monomers correlate with each other. The numerical prefactor in Eq. (3) comes from the spherical geometry of the problem and it could not be ignored. Note that, the Kuhn length and the second Virial coefficient could be calculated to a good precision numerically. About the PE chain in good solvent, Fig. (2) shows that the chain extension scales as ~ *N*^3/5^ as well. The difference between the PE and neutral chains has something to do with the correlations. The information regarding correlations is stored in the second Virial coefficient. The second Virial coefficient for the PE in good solvent for *l*_B_ = 0.7 σ is *b* = 2.99 σ^3^, for *l*_B_ = 2.0σ is *b* = 2.5 σ^3^ and for the PE in poor solvent for *l*_B_ = 0.7 σ is *b* = 0.01. These numerical values justify well why the PE extension is larger than the neutral chain in good solvent. Moreover, they tell us that the PE chain shrinks as the Bjerrum length increases. In poor solvent for *l*_B_ = 0.7, the second Virial coefficient almost vanishes. So one must consider the third Virial coefficient to tackle the PE problem in poor solvent. Note that considering the third Virial coefficient has no analytical solution. So I do not address that here.

**Figure 2:**
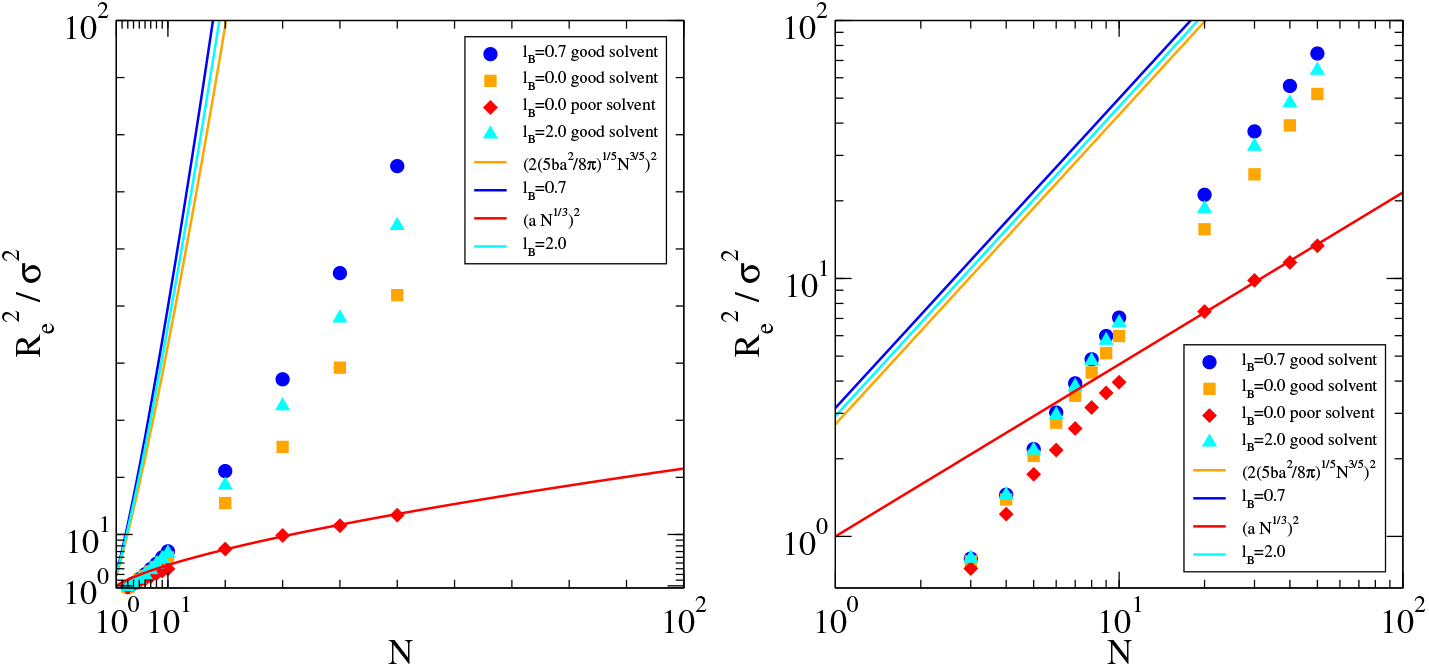
Both figures show the mean squared end-to-end distance as a function of the degree of polymerization. In the left figure, the linear plot and in the right figure the logarithmic plot is shown. The solid lines show the theories and the discrete symbols show the MD results.

## CONCLUSION

In this study, the PEs are investigated theoretically and numerically. The calculations show that in good solvent the PE chain extension scales with the degree of polymerization very much similar to the neutral chain. In this research I did not see any evidence that the PEs behave like ideal chains. The idea that has been mentioned in a the work already published (10) based on the blob picture. The DFT framework and the MD simulations predict a larger chain extension for the PE in good solvent while keeping the power laws untouched. Also, they predict a chain shrinkage upon increasing the Bjerrum length of solvent. Therefore, we see that implicit approach to the PE problem works well either through theory or simulation. The major outcome of this study is that in biological media the electrostatic interactions solely alter the correlations between monomers and do not touch the power laws. This suggests that the works already published must be probably reviewed.

## ACKNOWLEDGMENTS

I thank the Cold Spring Harbor Laboratory (CSHL) and the Regeneron Pharmaceuticals Inc. for financial support, Wolfram Research for Mathematica, the Canonical Ltd. for Ubuntu Linux operating system, the GNU Fortran compiler, the Weizmann Institute of Science for XMGrace, University of Illinois at Urbana-Champaign and the National Institute of Health (NIH) for the Visual Molecular Dynamics (VMD) and the Overleaf latex editor.

## Notes

### Competing Interest Statement

The authors have declared no competing interest.

## REFERENCES

1. Rubinstein, M., and R. Colby, 2004. Polymer Physics. OUP Oxford, Oxford, first edition.

2. Schwabl, F., and W. Brewer, 2006. Statistical Mechanics, Advanced Texts in Physics. Springer Berlin Heidelberg, Oxford, first edition.

3. Doi, M., and S. F. Edwards, 1986. Theory of polymer dynamics. Oxford science publication, Oxford, first edition.

4. Gennes, P. G., 1979. Scaling Concepts in Polymer Physics. Cornell University Press, Oxford, first edition.

5. Hirz, S. J., 1979. Modeling of Interactions Between Adsorbed Block Copolymers. University of Minnesota, Minnesota, first edition.

6. Safran, S., 2003. Statistical Thermodynamics Of Surfaces, Interfaces, And Membranes. Boca Raton: CRC Press., Oxford, first edition.

7. Debye, P., and E. Hüeckel, 1923. Zur Theorie der Elektrolyte. I. Gefrierpunktserniedrigung und verwandte Erscheinungen.

7. Smit, B., and D. Frenkel, 1996. Understanding Molecular Simulation: From Algorithms to Applications. Academic Press, Oxford, first edition.

7. Kremer, K., and G. S. Grest, 1990. Dynamics of entangled linear polymer melts: A molecular-dynamics simulation. The Journal of Chemical Physics 92.

7. Kreer, T., 2016. Polymer-brush lubrication: a review of recent theoretical advances. Soft Matter 12.

